# Irreversible PDMS bonding using flame activation of adhesives for fabrication of microfluidic and organ-on-chip devices

**DOI:** 10.1101/2023.08.17.553712

**Authors:** Ryan Singer, Jeremy A. Hirota, Mohammadhossein Dabaghi

**Author notes:** corresponding authors contact information: Ryan Singer –.

## Abstract

Polydimethylsiloxane (PDMS) is widely used for microfluidics fabrication in many disciplines due to its ease of use in soft lithography and its ability to bond liquid-tight seals. A variety of PDMS-PDMS bonding methods exist, but each may have limitations for some applications. For example, chemical bonding via oxygen plasma treatment is reliable but requires expensive equipment and specialized training. Here we present a rapid, low-cost, and accessible method for irreversible PDMS bonding in which flame treatment activates PDMS and Nitto 5302A adhesive surfaces. Using this technique, PDMS microchannels can be fabricated with a bonding integrity of up to 325 kPa burst pressure. This technique is suitable for fabricating organ-on-chip devices for cell line and primary cell culture, supporting cell viability and establishment of key cellular features in apical and basal compartments, while maintaining bonding integrity.

## 1. Introduction

The use of polydimethylsiloxane (PDMS) for fabricating microfluidic devices is becoming increasingly prominent in biomedical research due to its ability to form liquid-tight channels. Common methods for PDMS-PDMS bonding utilize the generation of reactive functional groups for covalent bonding, known as surface activation.[1]–[3] Several adhesive-based methods have also been developed for irreversibly bonding PDMS. [3]–[8] Adhesives are advantageous due to their accessibility, as well as enabling simple integration of membranes in microfluidic devices. Membrane integration is crucial in microfluidic applications that require separation or compartmentalization, such as analytical chemistry and organ-on-chip devices. In chemically bonded PDMS devices, the selection of membrane materials is limited to those which are chemically compatible, whereas adhesives enable liquid-tight sealing to a wide range of materials. [2], [5], [7], [9]

PDMS surface activation methods and several adhesives for PDMS-PDMS bonding have been independently characterized, but surface activation in conjunction with adhesives has not been demonstrated to our knowledge. We present a rapid, low-cost, and accessible method for high-strength irreversible PDMS bonding via flame activation of adhesives.

## 2. Materials and Methods

### 3.1. Adhesives

Four commercially available double-sided adhesives were evaluated for their ability to bond to PDMS following flame treatment: **Nitto 5302A** (Nitto Denko, 5302A-50); **ARclean® 90176** (Adhesives Research, AR-90176); **Scotch™ 7951** (3M, 7000050083); and **ARcare® 92712** (Adhesives Research, AR-MH-92712). The materials and thicknesses of each adhesive type are reported in Table 1.

**Table 1.**
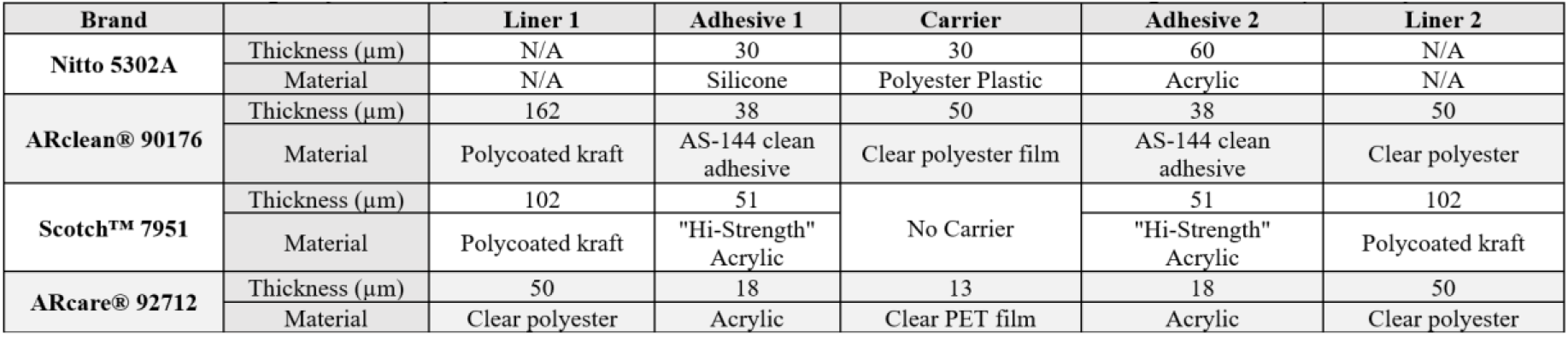
Product specification for adhesives used in this work based on the data provided by manufacturers.

### 3.2. Applying flame treatment

A small butane torch (Sondiko, Amazon, sondiko-1920) was used to activate the PDMS surface as described in previous work. [10] Briefly, the end of the torch nozzle was maintained at 2 cm from the surface (flame temperature ∼ 1300 ºC) and the flame was exposed for 5 s while sweeping the flame evenly at roughly 0.5 m/s. Following removal of the release liner, the adhesive surface was exposed using the same method but for only 3 s with a torch movement speed of roughly 1 m/s.

### 3.3. Microfluidic device fabrication

Photolithography molds were cast with PDMS (Dow SYLGARD™ 184, Ellsworth Adhesives Canada, 2085925) of 10:1 (wt:wt) base-to-curing agent ratio (**Figure 1AI-II and BI-II**). Adhesives were cut using a cutting machine (Silhouette Cameo® 3) before removing the release liner and flame-treating the adhesives and PDMS parts (**Figure 1AIII and BIII**). The activated adhesives were pressed firmly against the respective activated PDMS surfaces before removing the second release liners and firmly sandwiching a polyester membrane (Sterlitech, 1300018) between the apical and basal channels (**Figure 1AIV-V, BIV-V, and C**). All microfluidic devices were autoclaved prior to cell culture.

**Figure 1.**
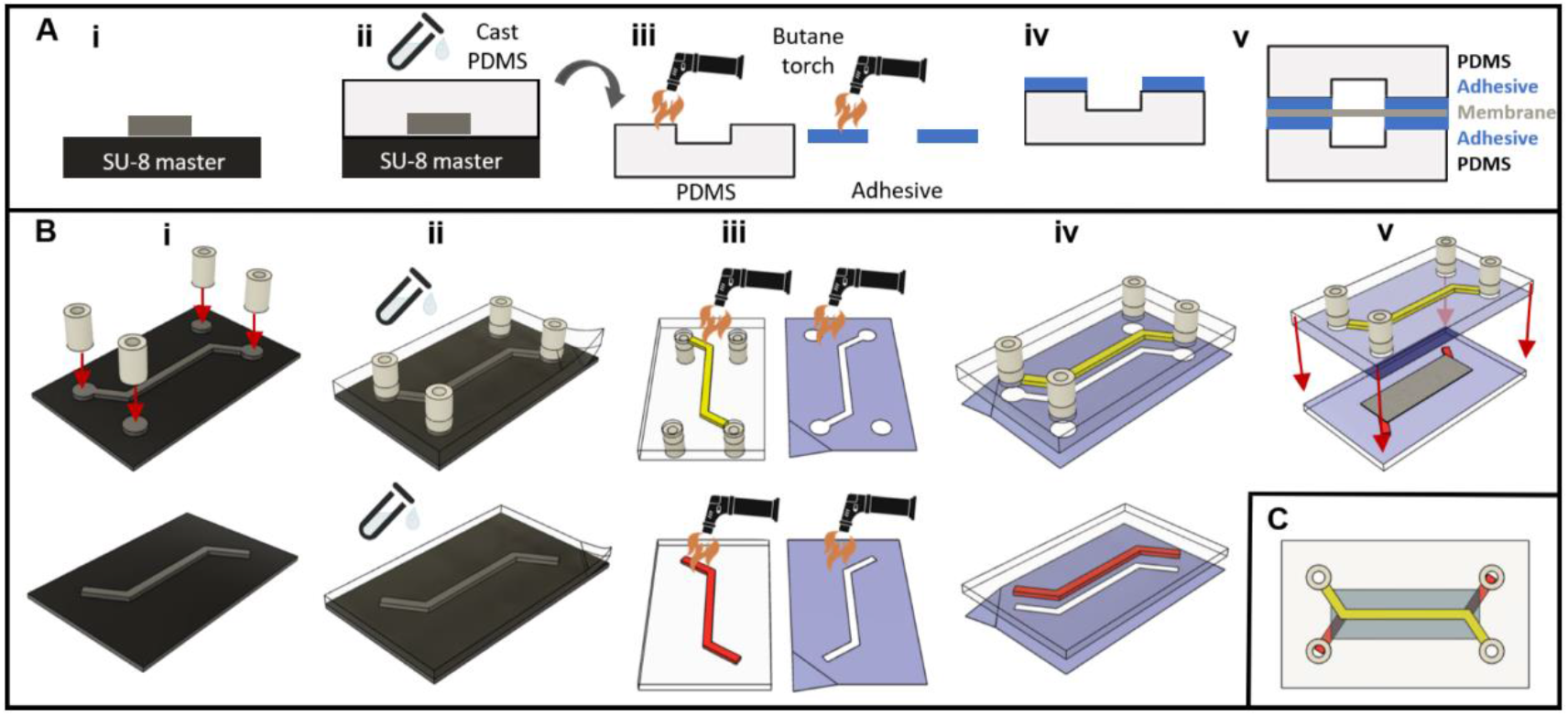
Fabrication of membrane-separated microchannels using flame-activated double-sided adhesives and PDMS. A) Cross-sectional and B) perspective depictions of the fabrication process. i) Apical and basal photolithography molds of microchannel geometry are prepared, and PTFE tubing is placed on inlet/outlet ports before (ii) PDMS is cast and removed. The PDMS and adhesive surfaces are (iv) flame-treated and (iv) pressed firmly together before (v) sandwiching a membrane between the apical and basal parts. C) Top-view depiction of an assembled device with coloured apical (yellow) and basal (red) channels.

### 3.4. Cell culture and fluorescence imaging

Calu-3 cells were expanded with Minimum Essential Medium Alpha (αMEM, Corning®, CA45000-300) supplemented with 10% fetal bovine serum (FBS, Wisent Inc., 080-450), 1% HEPES (Corning®, 25-060-CI), and 1% antibiotic/antimycotic solution (Gibco™, 15240062). The Calu-3 cells were cultured in the apical channels of pristine, O_2_ plasma-treated, and flame-treated Nitto 5302A adhesive-bonded devices with constant basal αMEM perfusion of 20 µL/min for 3 weeks. The barrier function of the Calu-3 monolayer was terminally evaluated via immunofluorescence imaging of tight junction protein 1 (ZO-1 antibody, Invitrogen, 339194) and permeability assays using 500 µL of 2 mg/mL FITC-dextran (4 kDa, Chondrex Inc., 4013) on the apical cell surface and measuring the absorbance of the basal media effluent after 15 h of incubation and perfusion.

Primary human lung fibroblast cells (HLFCs) were extracted from human lung tissue samples in accordance with Hamilton integrated Research Ethics Board (HiREB 5099T, 5305T). The HLFCs were expanded with Dulbecco’s Modified Eagle Medium (DMEM, Gibco™, 11965-118) supplemented with 10% FBS and 1% antibiotic/antimycotic solution. HLFCs were seeded in the basal channel of flame-treated Nitto 5302A adhesive-bonded devices. Establishment of the HLFC cytoskeleton was evaluated at confluency via immunofluorescence staining of actin filaments using phalloidin-iFluor 488 (Invitrogen™, ab176753).

Calu-3 and HLFC viability were assessed using the LIVE/DEAD™ cell imaging kit (Invitrogen™, R37601). All fluorescent images were acquired using an EVOS™ M7000 microscope (Invitrogen™, AMF7000HCA).

### 3.5. Water contact angle and burst pressure measurement

The static sessile drop method was used to measure the contact angles of all substrates using 2 µL DI water droplets. Burst pressures were measured using a syringe pump connected to a sealed microchannel filled with DI water and a pressure transducer (TruWave Transducer, Edwards Lifesciences LLC, Irvine, CA, USA).

## 3. Results and Discussion

### 4.1. Selecting an adhesive based on surface energy and bonding strength

Surface activation via oxygen plasma or flame treatment increases the surface energies of both acrylic- and silicone-based materials, rendering them more hydrophilic.[3], [10] The ideal adhesive for PDMS bonding should exhibit PDMS-like surface energy following surface activation. ARclean® 90176, Scotch™ 7951, and ARcare® 92712 contain acrylic adhesives on both sides, whereas Nitto 5302A contains a silicone-based adhesive opposite an acrylic adhesive. The water contact angle measurements in this work demonstrate that PDMS and each adhesive type were more hydrophilic with flame treatment compared to O_2_ plasma treatment (**Figure 2A**). The mean water contact angles following flame treatment for PDMS (1.552 ± 0.974º) and Nitto 5302A silicone-based adhesive (1.036 ± 0.644º) were most similar, indicating that they are most likely to form the strongest bond among the adhesive options. Burst pressure measurements confirmed that Nitto 5302A forms the strongest bond with PDMS (**Figure 2B**).

**Figure 2.**
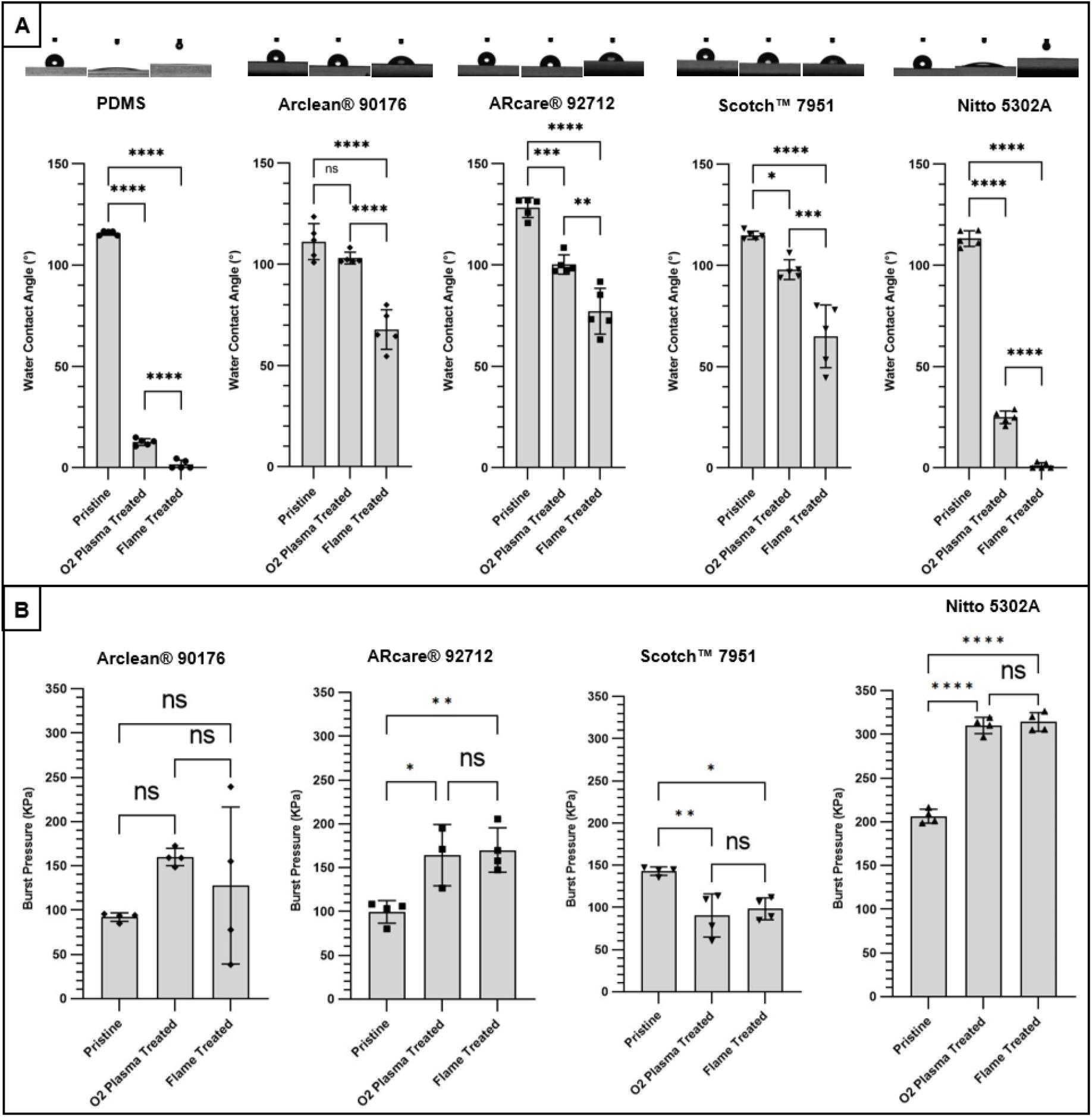
Nitto 5302A adhesive is most suitable for PDMS bonding. A) Water contact angle measurements for PDMS and each adhesive type under pristine (not treated), oxygen plasma-treated, and flame-treated conditions. B) Burst pressure measurements for PDMS microchannels bonded using pristine, oxygen plasma-treated, and flame-treated adhesives. Asterisks *, **, or **** indicate P ≤ 0.05, 0.01 or 0.0001, respectively.

### 4.2. Flame-treated Nitto 5302A adhesive supports sustained microfluidic cell cultures

Flame-treated Nitto 5302A devices maintained 325 kPa burst pressure following autoclaving and 3-week Calu-3 cell culture, whereas O_2_ plasma-treated Nitto 5302A devices withstood 25% less pressure (**Figure 2B and 3A**). These results were consistent with previous evidence that flame treatment provides higher quality surface activation of PDMS than oxygen plasma treatment.[10] Flame-treated Nitto 5302A devices could support high viability of Calu-3 cells for 3 weeks (**Figure 3B and C**). Immunofluorescent imaging of ZO-1 revealed ubiquitous formation of tight junctions throughout the apical channel (**Figure 3D)** capable of inhibiting diffusion of 4 kDa dextran (**Figure 3E**). Flame-treated Nitto 5302A devices were also capable of supporting a high viability of HLFCs in basal channels (**Figure 3F and G**) and the establishment of actin filament cytoskeleton networks (**Figure 3H**).

**Figure 3.**
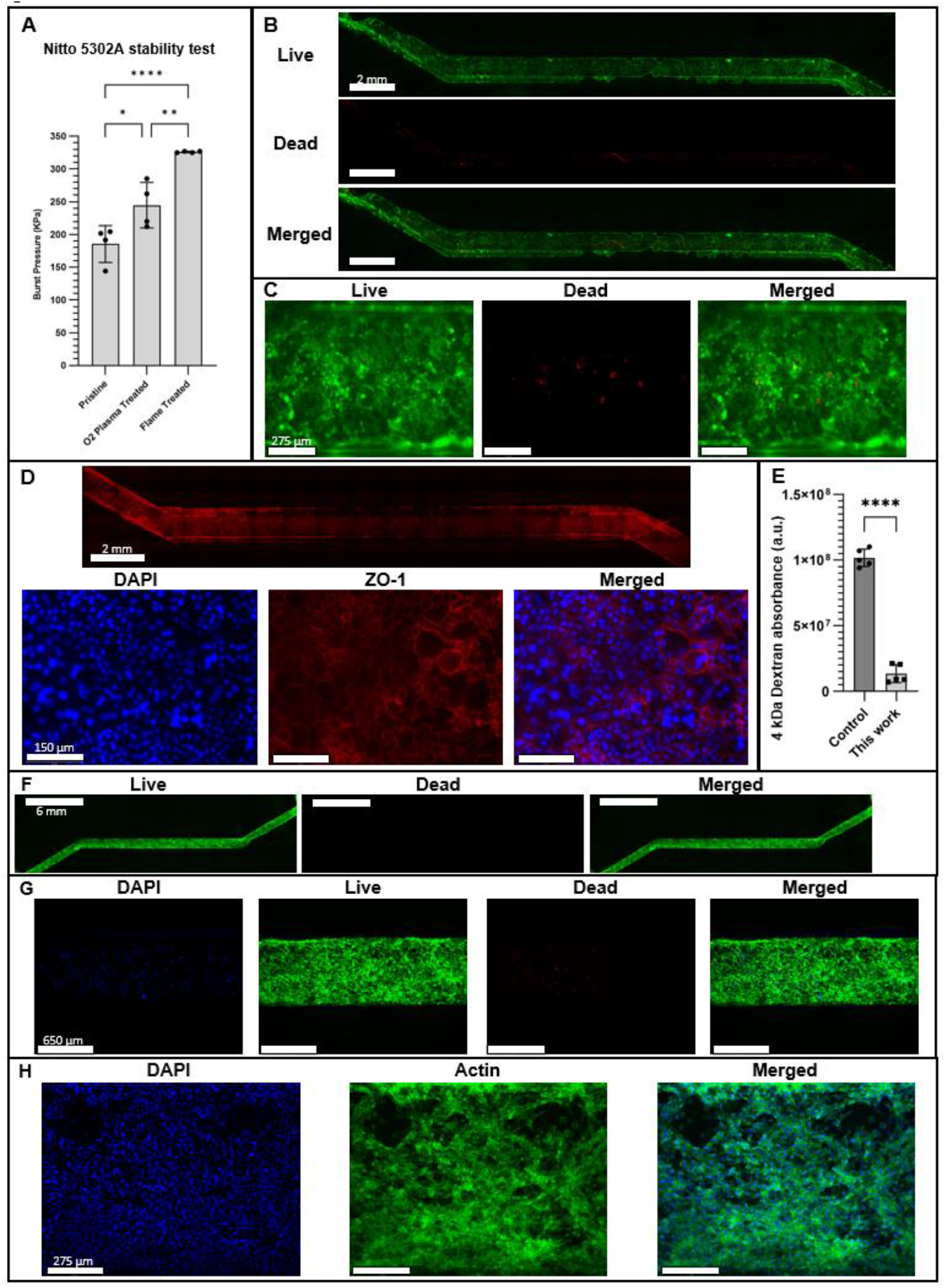
Flame-treated Nitto 5302A adhesive-bonded organ-on-a-chip devices are suitable for sustained cell culture. A) Burst pressure measurements for organ-on-chip devices bonded using pristine, oxygen plasma-treated, and flame-treated Nitto 5302A adhesive following 3-week Calu-3 culture. B) Full channel and (C) magnified live/dead fluorescence images of Calu-3 cells following 3-week culture. D) Full channel and magnified ZO-1 (red) and cell nuclei (blue) fluorescence images. E) Basal effluent absorbance of apically diffused 4 kDa dextran before (control) and after 3-week Calu-3 culture. F) Full channel and (G) magnified live/dead and (H) actin fluorescence images of HLFCs following 2-week culture. Asterisks *, **, or **** indicate P ≤ 0.05, 0.01 or 0.0001, respectively.

## 4. Conclusions

A rapid and accessible method for irreversibly bonding PDMS has been presented. By flame-treating Nitto 5302A adhesive and PDMS, microchannels can be fabricated with a bonding strength of up to 325 kPa burst pressure. This technique is suitable for fabricating organ-on-chip devices with strong bonding integrity without capital investment for dedicated equipment, that can support sustained viability and establishment of key cellular features in Calu-3 epithelial cells and HLFCs.

## References

[1] A. Shakeri, S. Khan, and T. F. Didar, “Conventional and emerging strategies for the fabrication and functionalization of PDMS-based microfluidic devices,” Lab Chip, vol. 21, no. 16, pp. 353–375, 2021, doi: 10.1039/d1lc00288k.

[2] R. Sivakumar and N. Y. Lee, “Microfluidic device fabrication mediated by surface chemical bonding,” Analyst (London), vol. 145, no. 12, pp. 496–411, 2020, doi: 10.1039/d0an00614a.

[3] A. Borók, K. Laboda, and A. Bonyár, “PDMS bonding technologies for microfluidic applications: A review,” Biosensors, vol. 11, no. 8. MDPI, Aug. 01, 2021. doi: 10.3390/bios11080292.

[4] M. Chu, T. T. Nguyen, E. K. Lee, J. L. Morival, and M. Khine, “Plasma free reversible and irreversible microfluidic bonding,” Lab Chip, vol. 17, no. 2, pp. 267–273, Jan. 2017, doi: 10.1039/C6LC01338D.

[5] S. Hassanpour-Tamrin, A. Sanati-Nezhad, and A. Sen, “A simple and low-cost approach for irreversible bonding of polymethylmethacrylate and polydimethylsiloxane at room temperature for high-pressure hybrid microfluidics,” Sci Rep, vol. 11, no. 1, Dec. 2021, doi: 10.1038/s41598-021-83011-8.

[6] C. S. Thompson and A. R. Abate, “Adhesive-based bonding technique for PDMS microfluidic devices,” Lab Chip, vol. 13, no. 4, pp. 632–635, Feb. 2013, doi: 10.1039/c2lc40978j.

[7] M. Dabaghi et al., “Adhesive-Based Fabrication Technique for Culture of Lung Airway Epithelial Cells with Applications in Cell Patterning and Microfluidics,” ACS Biomater Sci Eng, vol. 7, no. 11, pp. 5301–5314, Nov. 2021, doi: 10.1021/acsbiomaterials.1c01200.

[8] S. Satyanarayana, R. N. Karnik, and A. Majumdar, “Stamp-and-stick room-temperature bonding technique for microdevices,” Journal of microelectromechanical systems, vol. 14, no. 2, pp. 392–399, 2005, doi: 10.1109/JMEMS.2004.839334.

[9] J. de Jong, R. G. H. Lammertink, and M. Wessling, “Membranes and microfluidics: a review,” Lab Chip, vol. 6, no. 9, p. 1125, 2006, doi: 10.1039/b603275c.

[10] R. Ghaemi, M. Dabaghi, R. Attalla, A. Shahid, H. H. Hsu, and P. R. Selvaganapathy, “Use of flame activation of surfaces to bond PDMS to variety of substrates for fabrication of multimaterial microchannels,” Journal of Micromechanics and Microengineering, vol. 28, no. 8, May 2018, doi: 10.1088/1361-6439/aabd29.

